# Host age-dependent evolution of a plant RNA virus

**DOI:** 10.1101/2022.06.28.497762

**Authors:** Izan Melero, Rubén González, Santiago F. Elena

**Affiliations:** Instituto de Biología Integrativa de Sistemas (CSIC - Universitat de València), Paterna, 46182 València, Spain; The Santa Fe Institute, Santa Fe NM 87501, USA

**Keywords:** experimental evolution, plant-virus interaction, developmental stages, virus evolution

## Abstract

Viruses are obligate pathogens that entirely rely on their host resources to complete their infectious cycle. The availability of such resources depends upon external and internal factors, being host age one of the most relevant ones. The interplay between host age and virus evolution has not been thoroughly studied in plants. Here, we have used the *Arabidopsis thaliana* - turnip mosaic virus (TuMV) pathosystem to study plant-virus interactions and virus evolution at three different host ages: vegetative (juvenile), bolting (transition) and reproductive (mature) stages. After infecting plants with a naïve and a well-adapted TuMV isolates, we observed that the older the host the faster and more severe the infections were. The same trend was observed for several other viruses. Thereafter, we experimentally evolved lineages of the naïve and the well-adapted TuMV isolates in plants from each of the three developmental stages. All evolved viruses enhanced their infection phenotypes, being this increase more intense on viruses evolved in younger hosts. The genomic changes of the evolved viral lineages revealed mutation patterns that strongly depended on the founder viral isolate as well as on the age of the host wherein the lineages evolved.

## Introduction

Viruses are obligate intracellular pathogens that hijack their hosts’ cellular components. When a virus takes over the control of host cells gene expression, cellular resources are diverted to generate viral factories and start producing all required viral components, ending up in the assembly and liberation of progeny virions. Because of that, the biochemical and physiological status of host cells will strongly impact the ability of a virus to accomplish its replication. This cellular status will depend on multiple and diverse factors: from the host genetics to the environmental components the host faces [1]. These internal biotic and external abiotic conditions drive the host-virus interactions, meaning that different host conditions may result in different susceptibilities to infection. Host homogeneity is rare in natural populations, which implies that viruses must face host populations whose individuals are genetically diverse and have different degrees of susceptibility to infection [2,3]. The interaction between pathogens and hosts with genetic variability in susceptibility to infection has been well studied [3-5], as well as the influence of developmental stages on host’s resistance to pathogens [6-9].

Variation in disease susceptibility among hosts of different age or life stages constitute one of the most important components of host heterogeneity affecting the outcome of host-pathogen interactions and their epidemiological dynamics [10]. These differences in susceptibility and resistance might imply that a virus adapted to preferentially infect a given age class, might be unsuccessful infecting other age classes. In nature, an interaction between host age and susceptibility to infection has been observed in many animal and plant systems for a wide range of pathogens such as bacteria, fungi, or viruses. In animals, young individuals are more susceptible to infection than older individuals for many viruses [11-13]. However, for other pathogens such as fungi, older animals may exhibit increased susceptibility compared to juvenile ones [14]. In plants, a common observation is that hosts become more resistant to pathogens as they mature [15-17]. This is explained by the existence of a plant mechanism known as age related resistance (ARR), which was first described in the *Arabidopsis thaliana* - *Pseudomonas syringae* pathosystem [18]. ARR is a plant defense response characterized by an increased resistance to pathogens in mature plants compared to young plants. However, some contradictory results on the role of ARR have been brought forward. *E*.*g*., onion plants can be more susceptible to certain fungal pathogens as they age [19], older arabidopsis plants are more susceptible to some RNA viruses [20], and the aging of petunia plants correlates with an activation of the pararetrovirus petunia vein clearing virus in response to changes in DNA methylation patterns [21]. Therefore, even though ARR seems to be an extended phenomenon in plants, it is not universal and depends on particular pathosystems.

Due to their sessile lifestyle, plants need to finely coordinate their growth and development in order to optimize fitness through rapid and appropriate responses to the different stresses they might face. As a matter of fact, the life cycle of flowering plants can be considered as a succession of distinct growth phases. One of these phases is the transition from a juvenile vegetative to a mature reproductive stage. The transition to flowering is under the control of a complex genetic network that integrates information from both endogenous and environmental cues [22] where plants change their genetic programs to switch from growth to reproduction. Despite flowering not being the developmental transition needed for increased pathogen resistance [23], multiple studies have shown an interaction between flowering and defense. Resource allocation theories claim that plants have a limited pool of resources which they must employ to carry out different functions, therefore promoting trade-offs that determine resource allocation [24,25]. Among these trade-offs, a particularly well-known one is the reallocation of resources from defense to flower development. This reallocation makes sense for plants that reproduce only once in their lifespan and need to produce descendants at any cost. Interestingly, infection with certain pathogens can speed up flowering [26,27]. Furthermore, it has been shown that pathogens face different defense responses depending on the developmental stage of the plant [16,28]. Age-specific modifications, such as the accumulation of reactive oxygen species or the reprogramming of hormone crosstalk pathways [29,30] influence the host susceptibility and, therefore, entail changing scenarios for pathogens. However, how these differences may affect virus evolutionary strategies remains unknown.

Although the influence of aging over pathogen infection has gathered the attention of many studies, less attention has been paid to whether different host ages represent different selective environments for viruses thus modulating their evolution. It is known that selection is expected to favor combinations of traits that maximize pathogen’s fitness. Most pathogen eco-evolutionary theories assume a link between virulence and transmission [31], as there should be a compromise between the duration of an infection (and therefore the time in which a pathogen can propagate and infect other hosts), and the damage that the pathogen causes to its host. Interestingly, it has been suggested that the virulence-transmission trade-off is affected by the host’s life history [9,32,33]. Even though parasite-induced host mortality is the most used measure of virulence for horizontally transmitted parasites [34], there are pathogens that follow other evolutionary strategies to guarantee their reproduction and transmission without prematurely killing the host and, in consequence, themselves. This is the case of castrating parasites, which aim to diminish host fitness through interference on host fecundity. In fact, host castration has been argued to be an explicit evolutionary strategy of the parasite [35-37], because by castrating their hosts, pathogens can redirect host resources that were previously allocated for reproduction to secure their own reproduction and survival. Because of the reallocation of resources along development [38], even if viruses are infecting the same host genotype, the selective pressures imposed by the host may differ depending on the host age. And therefore, adaptive mutations improving viral fitness in one host age may be selected against, or be neutral, in a different one.

Here we sought to better understand how host age influences virus evolution. We used *A. thaliana* (L.) Heynh, a model organism in plant research [39], to evaluate the susceptibility to virus infection along three developmental stages: (*i*) vegetative juvenile stage, where plants allocate resources to increase their size and mass, (*ii*) bolting, an indicator of developmental stage transition [40], and (*iii*) flowering, where mature plants allocate resources to reproduction. We evaluated the host-age effect on the susceptibility to a collection of viruses from six different genera. After characterizing the host-age dependent susceptibility, we performed an evolution experiment using turnip mosaic virus (TuMV; species *Turnip mosaic virus*, genus *Potyvirus*, family *Potyviridae*) [41]. TuMV has a 9.5 Kb positive single-stranded RNA genome that translates into a large polyprotein then processed by three viral-encoded proteases into ten mature products, plus a frameshift protein [42]. TuMV infects over 300 plant species, though most belong to the *Brassicaceae* family [41-43]. We used two different TuMV isolates: an isolate *naïve* to arabidopsis (hereafter referred as AS) and an experimentally preadapted one (referred as DV). The resulting evolved lineages were phenotyped and sequenced to study the effects of host age on virus evolution. In summary, this work aims to (*i*) measure the impact of arabidopsis age on susceptibility to virus infection, and (*ii*) evaluate how virus evolution is affected by its host age.

## Material and methods

### Plant material and growth conditions

All experiments were performed in a growing chamber at 24 °C during light time and 20 °C during dark, 45% relative humidity, and 125 µmol m^−2^ s^−1^ of light intensity (1:3 mixture of 450 nm blue and 670 nm purple LEDs). The photoperiod consisted in 16 h light/8 h dark for long-day conditions and 8 h light/16 h of dark for short-day ones.

Arabidopsis plants of the Col-0 accession were used as hosts. Plants were inoculated at three different developmental stages: prebolting (juvenile), bolting (transition) and postbolting (mature) (Fig. 1A). In long day conditions these stages correspond with plants with an age of 18, 25 or 32 days after sowing (das), respectively. For short day conditions each stage corresponds to 52, 66 or 73 das, respectively. As bolting can be used as indicator of the vegetative to reproductive phase change [40], each stage could be associated with different plant phases: vegetative growth, phase transition, and reproductive growth. Following the principal growth stages described by Boyes et al. [44], the inoculated plants also correspond to different principal growth stages: prebolting to stage 1, bolting to stage 3 and postbolting to stage 5.

**Figure 1.**
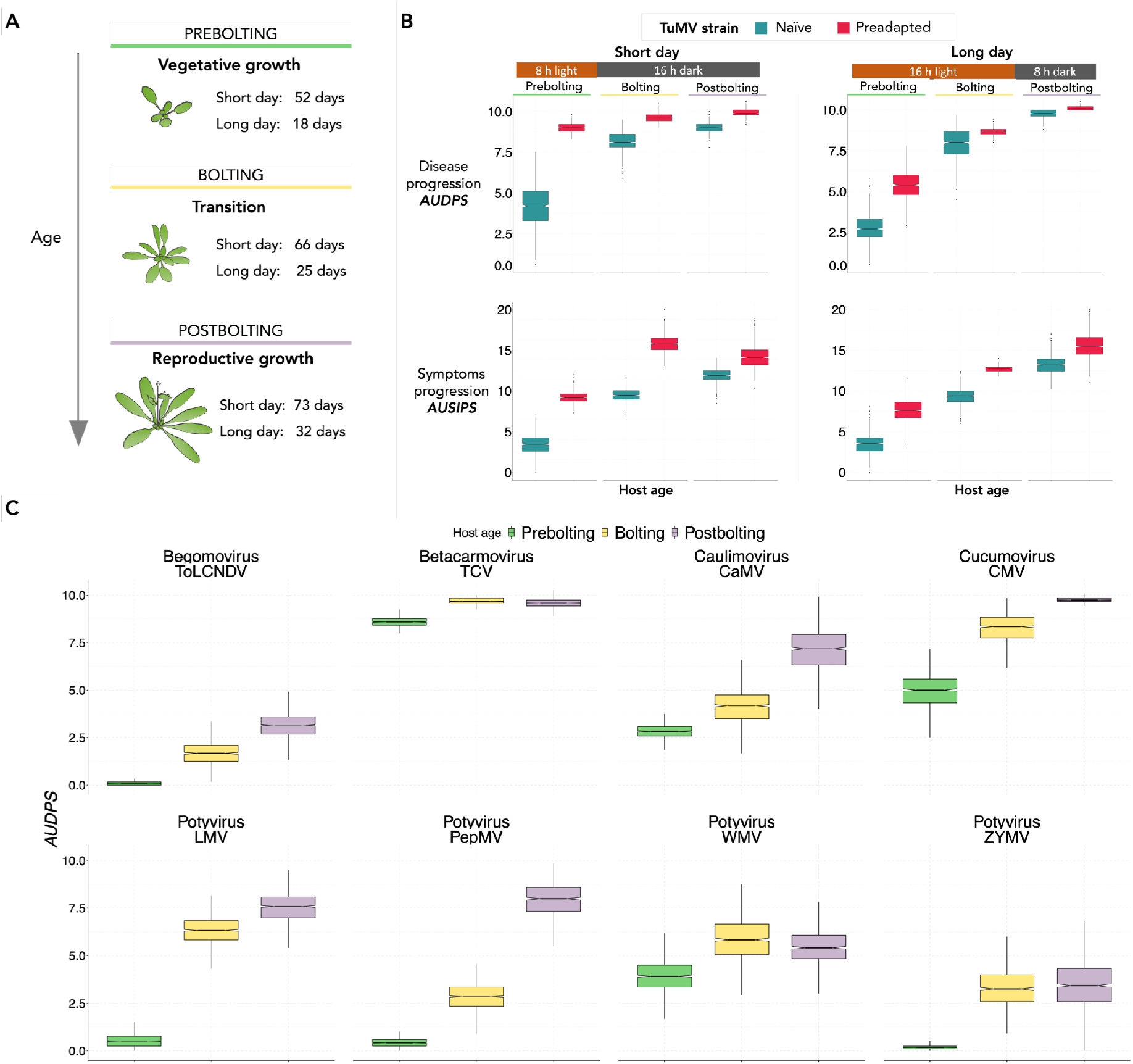
A) Host ages evaluated in this work and the corresponding number of days after sowing at which plants were inoculated at both photoperiod conditions. B) Infection phenotypes for TuMV in short- and long-day photoperiod conditions. Infectivity progression (*AUDPS* values; upper row) and symptomatology progression (*AUSIPS* values; lower row) for naïve AS (blue) and preadapted DV (red) TuMV in the three plant developmental stages evaluated. C) Infectivity progression (*AUDPS* values) for a set of viruses in the three plant developmental stages evaluated. For each virus its genus is indicated.

### Viruses and experimental evolution

Inoculations were done using homogenized virus-infected tissue preserved at −80 °C. The virus inoculum consisted of 100 mg of homogeneous N_2_-frozen infected tissue mixed with 1 mL of phosphate buffer and 10% Carborundum (100 mg/mL). For each virus and developmental stage condition, 12 plants were inoculated.

The following viruses have been used: *Begomovirus* (tomato leaf curl New Delhi virus, ToLCNDV), *Betacarmovirus* (turnip crinkle virus, TCV), *Caulimovirus* (cauliflower mosaic virus, CaMV), *Cucumovirus* (cucumber mosaic virus, CMV), *Potexvirus* (pepino mosaic virus, PepMV), and *Potyvirus* (lettuce mosaic virus, LMV; turnip mosaic virus, TuMV; watermelon mosaic virus, WMV; and zucchini yellow mosaic virus, ZYMV). Virus stocks were generated by harvesting and homogenizing infected tissue of arabidopsis (CaMV, TCV and TuMV-DV), *Nicotiana benthamiana* Domin (CMV, LMV, PepMV, TuMV-YC5, and WMV) or *Chenopodium quinoa* Willd (ZYMV and ToLCNDV). The two isolates of TuMV used differ in their degree of adaptation and severity of symptoms in arabidopsis. The naïve AS isolate came from strain YC5 (GenBank, AF53055.2) originally obtained from calla lily (*Zantedeschia sp*.), which was cloned under the 35S promoter and NOS terminator, resulting in the p35STunos infectious clone [45]. The arabidopsis-adapted DV isolate was obtained after experimentally evolving the AS isolate for twelve passages in Col-0 plants [3].

For the evolution experiment, 10 plants were inoculated per combination of developmental stage and TuMV strain. Fourteen days post-inoculation (dpi) the symptomatic infected plants were collected, making a pool of infected tissue that was homogenized and used as inoculum to start a five-passages evolution. For each one of the three developmental stages, three independent lineages were established to serve as biological replicates of the evolutionary process (Fig. 2).

**Figure 2.**
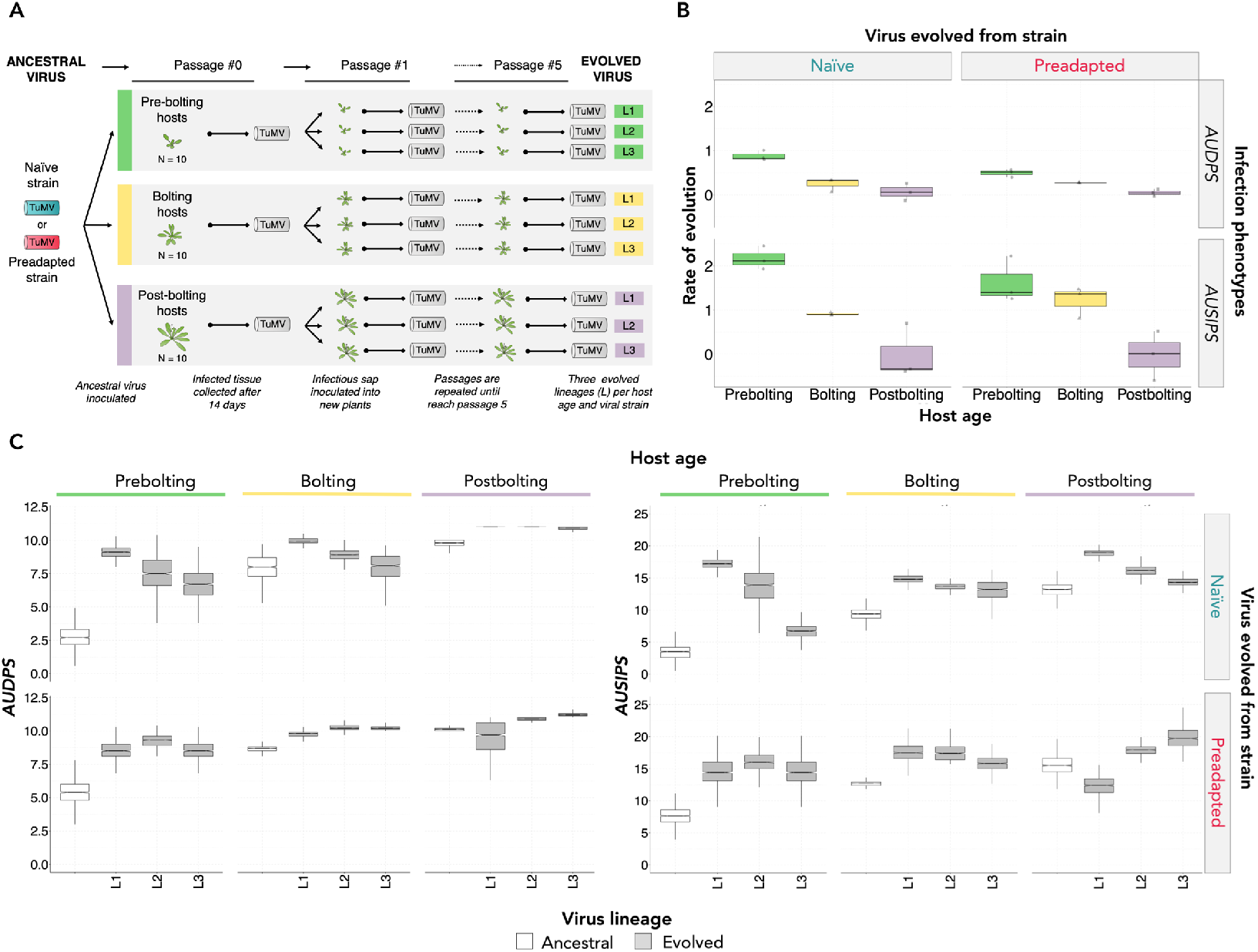
A) Experimental evolution design. B) Rates of phenotypic evolution for lineages evolved from the naïve AS isolate (left panel) and the preadapted DV one (right panel). Rates are calculated both for *AUDPS* (upper row) and *AUSIPS* (lower row). C) Evolved (grey) lineages infection phenotypes (*AUDPS* on the left; *AUSIPS* on the right) compared to their corresponding ancestral (white) viruses. Upper row shows the relative phenotype of viruses evolved from the naïve AS isolate, while the lower row represents the values for viruses evolved from the preadapted DV isolate.

### Infection characterization

Upon inoculation, plants were daily inspected for visual symptoms for fourteen days and annotated following a discrete scale of symptoms severity from absence of symptoms (0) to full necrosis of the plant (5) (as shown in Fig. 1 in [46]). The infectivity and severity of symptoms data along 14 dpi were used to calculate the area under the disease progress stairs (*AUDPS)* and intensity progression step (*AUSIPS)* values, respectively, as described in [47]. *AUDPS* and *AUSIPS* values were computed using the agricolae R package version 1.3-2 with R version 3.6.1 in RStudio version 1.2.1335.

### Statistical analyses

A bootstrapping method described in [47] was used to estimate the confidence intervals (95% CIs) of *AUDPS* and *AUSIPS*.

*AUDPS* and *AUSIPS* evolution data were fitted to a generalized linear model (GLM) that incorporated a Normal probability distribution and an identity link function. The model equation incorporated the developmental stage (vegetative, bolting and reproductive) and the virus isolate (AS and DV) as orthogonal factors, viral lineage as a nested factor within the interaction of these two factors, and passage number as a covariable. The significance of each factor was evaluated using a likelihood-ratio test that asymptotically follows a *χ*^2^ distribution. The magnitude of effects associated to each factor was evaluated using the 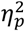 statistic (proportion of total variability in the variables attributable to each factor in the model; conventionally, values of 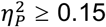 are considered as large effects). These analyses were performed with SPSS version 28.0.1.0 (IBM, Armonk, NY).

Rates of phenotypic evolution were evaluated by fitting the time-series data of *AUDPS* and *AUSIPS* to a first-order autoregressive integrated moving-average, ARIMA(1,0,0), model. The model equation fitted has the form:

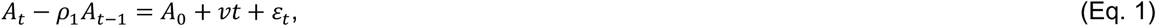

where *A*_*k*_ represents the disease phenotypic values at passage *k, ρ*_1_ measures the degree of self-similarity in the time-series data (correlation between values at passages *t* and *t* – 1), *ε*_*t*_ represents the sampling error at passage *t*, and *v* represents the dependency of *A* with passage number, that is, the rate of phenotypic evolution.

All statistical analyses were done with R version 3.6.1 in RStudio version 1.2.1335 unless otherwise indicated.

### TuMV genomes sequencing

RNA was extracted from infected plant tissue using NZY Total RNA Isolation Kit (NZYTech, Portugal) following the described protocol and using a 30 mg quantity of plant material for extraction. The quality of the RNAs used to prepare RNA-seq libraries was checked with the Qubit RNA BR Assay Kit (Thermo Fisher, USA). SMAT libraries, Illumina sequencing (paired end, 150 bp), and quality check of the RNA-seq libraries were done by Novogene Europe. Nineteen bases from the 5′ end and 12 from the 3′ of the reads were trimmed with cutadapt version 2.10 [48]. Trimmed sequences were mapped with HiSat2 version 2.1.0 [49] to the YC5 genome with a modified minimum score parameter (L, 0.0, −0.8) to allow more mismatches. Resulting SAM files were BAM-converted, sorted, indexed, and analyzed with SAMtools version 1.10 [50]. SNP calling was performed using bcftools version 1.6 by first using the mpileup subroutine.

## Results

### Host aging increases susceptibility to TuMV infection independently of the viral isolate or the photoperiod conditions

To test the effect of host developmental stage on disease traits (*AUDPS* and *AUSIPS*), we inoculated the arabidopsis-naïve AS and the arabidopsis-adapted DV isolates [3] in plants at the three afore mentioned developmental stages. Firstly, we observed that *AUDPS* values were significantly different between host’s developmental stages, regardless the viral genotype and photoperiod conditions (Fig. 1B). In all cases, disease progressed faster as plants became older (Fig. 1B, upper row). For instance, in long day conditions, the naïve AS isolate showed the lowest *AUDPS* in prebolting plants (mean ±1 SD; 2.688 ±0.015, which increased in bolting plants (7.930 ±0.043) and further increased in postbolting plants (9.783 ±0.053). The same trend was observed for plants infected with the preadapted DV isolate: this virus infected postbolting plants (10.112 ±0.055) significantly better than plants at prebolting (5.331 ±0.0289) and bolting (8.672 ±0.047) stages. Moreover, as these results put in evidence, the preadapted DV virus infected significantly better than the naïve one in all developmental stages. Previous work [3] has described that, for TuMV inoculated in prebolting Col-0 plants at long-day conditions, infectivity progression correlates with virus accumulation and the severity of symptoms. Our results reinforce this observation, since the *AUSIPS* values completely mimic the results described for *AUDPS* (Fig. 1B, lower row).

Secondly, we tested the effect of the photoperiod conditions in the outcome of the experiment. For each photoperiod condition, we inoculated plants at the corresponding days after sowing until reaching all three of the developmental stages evaluated (see Material and methods for details). For short-day conditions, we observed the same trend described in the previous paragraph for long-day conditions: disease and symptomology progression were faster as plants aged (Fig. 1B). Hence, we rule out a possible effect of photoperiod in the observed results.

### The influence of aging over susceptibility to infection seems widespread among viruses

After showing the effect of aging in TuMV disease progression and severity, we sought to confirm whether this was a general feature or it was dependent on the viral species being studied. To this end, we inoculated a total of eight viruses belonging to five different genera: (see Material and methods for details). In these experiments, only *AUDPS* was evaluated. Overall, the results are reproducing those observed for TuMV: the older the arabidopsis plants, the more susceptible they were to infection (Fig. 1C). It is worth noticing that for some viruses, bolting and postbolting plants were significantly more susceptible than juvenile prebolting plants, but no significant differences existed between bolting and mature postbolting infected plants (Fig. 1C). This was the case for TCV (bolting, 9.668 ±0.177; postbolting, 9.571 ±0.241), WMV (bolting, 5.800 ±1.041; postbolting, 5.374 ±0.922) and for ZYMV (bolting, 3.275 ±0.990; postbolting, 3.497 ±1.295).

### Host age and virus past evolutionary history shape the evolution of disease phenotypes

After studying the role of host age on susceptibility to virus infection, we used the two aforementioned TuMV isolates to start a five-passages evolution experiment. Each isolate was evolved in arabidopsis plants at each developmental stage maintained under long-day conditions (Fig. 2A). The estimates of the two disease-related traits along evolutionary passages were fitted to the GLM described in the Materials and methods section. For both, developmental stage had a highly significant and large effect (*AUDPS*: *χ*^2^ = 325.330, 2 d.f., *P* < 0.001, 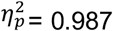 *AUSIPS*: *χ*^2^ = 148.497, 2 d.f., *P* < 0.001, 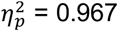), with mature plants having larger values for both variables, followed by bolting and prebolting. Likewise, the viral isolate also exerted a major and highly significant in both traits consistent along the evolution experiment (*AUDPS*: *χ*^2^ = 22.679, 1 d.f., *P* < 0.001, 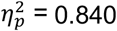 *AUSIPS*: *χ*^2^ = 9.133, 1 d.f., *P* = 0.003,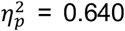), with DV being, overall, more virulent than AS. Moreover, these effects were not independent to each other: a significant and of large magnitude interaction between both factors was also observed for *AUDPS* (*χ*^2^ = 17.481, 2 d.f., *P* < 0.001,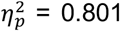) but not for *AUSIPS* (*χ*^2^ = 3.772, 2 d.f., *P* = 0.152). More relevant, the two disease variables significantly increased in value during the five-passages evolution experiment, being the overall effect of evolutionary time also of large magnitude (*AUDPS*: *χ*^2^ = 91.138, 1 d.f., *P* < 0.001, 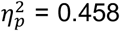 *AUSIPS*: *χ*^2^ = 120.699, 1 d.f., *P* < 0.001, 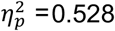). Finally, the effect of evolutionary time was affected by the developmental stage of the plants (*AUDPS*: *χ*^2^ = 49.005, 2 d.f., *P* < 0.001, 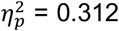 *AUSIPS*: *χ*^2^ = 62.622, 2 d.f., *P* = 0.191, 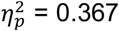), being larger for juvenile plants than for mature ones. No interaction between evolutionary time and viral isolate was detected (*i*.*e*., ancestral differences were maintained along the experiment), nor higher-order interactions or significant differences among independent lineages within each treatment.

The *AUDPS* and *AUSIPS* values of the evolved viruses were compared to the values obtained for their corresponding ancestral (Fig. 2C). We observed that the disease and symptomatology progression were faster in all the evolved viruses in comparison with their antecessors. However, viruses evolved in juvenile (prebolting) hosts experienced larger increases in disease severity (95% CI of ancestral viruses *vs*. 95% CI of evolved viruses), and this increase was even larger for the naïve AS isolate ([2.293 −2.410] *vs*. [7.678 - 7.753]) than for the preadapted DV one ([5.766 - 5.877] *vs*. [8.226 - 8.283]), when comparing with bolting ([7.873 - 7.978] *vs*. [8.642 - 8.699]; [8.690 – 8.721] *vs*. [9.863 – 9.884]) or postbolting hosts ([9.684 - 9.711] *vs*. [10.588 - 10.633]; [10.094 - 10.106] *vs*. [10.558 – 10.601]).

At the end of the evolution experiment, the rate of phenotypic evolution for the two disease-related traits *AUDPS* and *AUSIPS* was evaluated using the ARIMA model shown in Eq. 1. Our results show that the younger the host, the faster the evolution of both disease traits (Fig. 2B). This happened on the naïve AS isolate, both for disease progression *AUDPS*, where the evolutionary rate was higher in prebolting (average across lineages: 0.881) than in bolting (0.244) and postbolting (0.066) plants; and for symptomatology progression *AUSIPS*: the evolutionary rate was higher in prebolting (2.166) than in bolting (0.912) and postbolting (−0.012) plants. A qualitatively similar pattern was observed for the preadapted DV isolate: the rate of *AUDPS* evolution was higher in prebolting (0.496) than in bolting (0.276) and postbolting (0.048) plants; and for *AUSIPS* was also higher on prebolting (1.626) than in bolting (1.216) and postbolting (−0.061) plants. These results are consistent with those obtained with the GLM analysis.

### Genomic changes in evolved viruses dependent on the ancestral viral isolate and the host age

The genomes of the evolved viruses were sequenced to identify mutations that arose during the experimental evolution process (Fig. 3). All lineages evolved from the naïve AS isolate had the same non-synonymous mutation in the VPg/N2039D, regardless of the developmental stage of the host in which they were evolved. Interestingly, this mutation affects the same amino acid residue that mutation VPg/N2039H that was already present in the preadapted DV isolate. Therefore, mutations at position 2039 of VPg might be involved into adaptation to arabidopsis. However, these two mutations likely have strongly different effects in VPg structure and function: whereas N2039D replaces a polar amino acid by a negatively charged residue, N2039H replacement implies the presence of a bulky and positively charged residue. Other nonsynonymous mutations affecting HC-Pro, CI or NIb have been observed in other lineages, yet none was shared. By contrast, the three AS-derived lineages evolved in prebolting hosts, shared a nonsynonymous mutation in CI (amino acid residue 1468).

**Figure 3.**
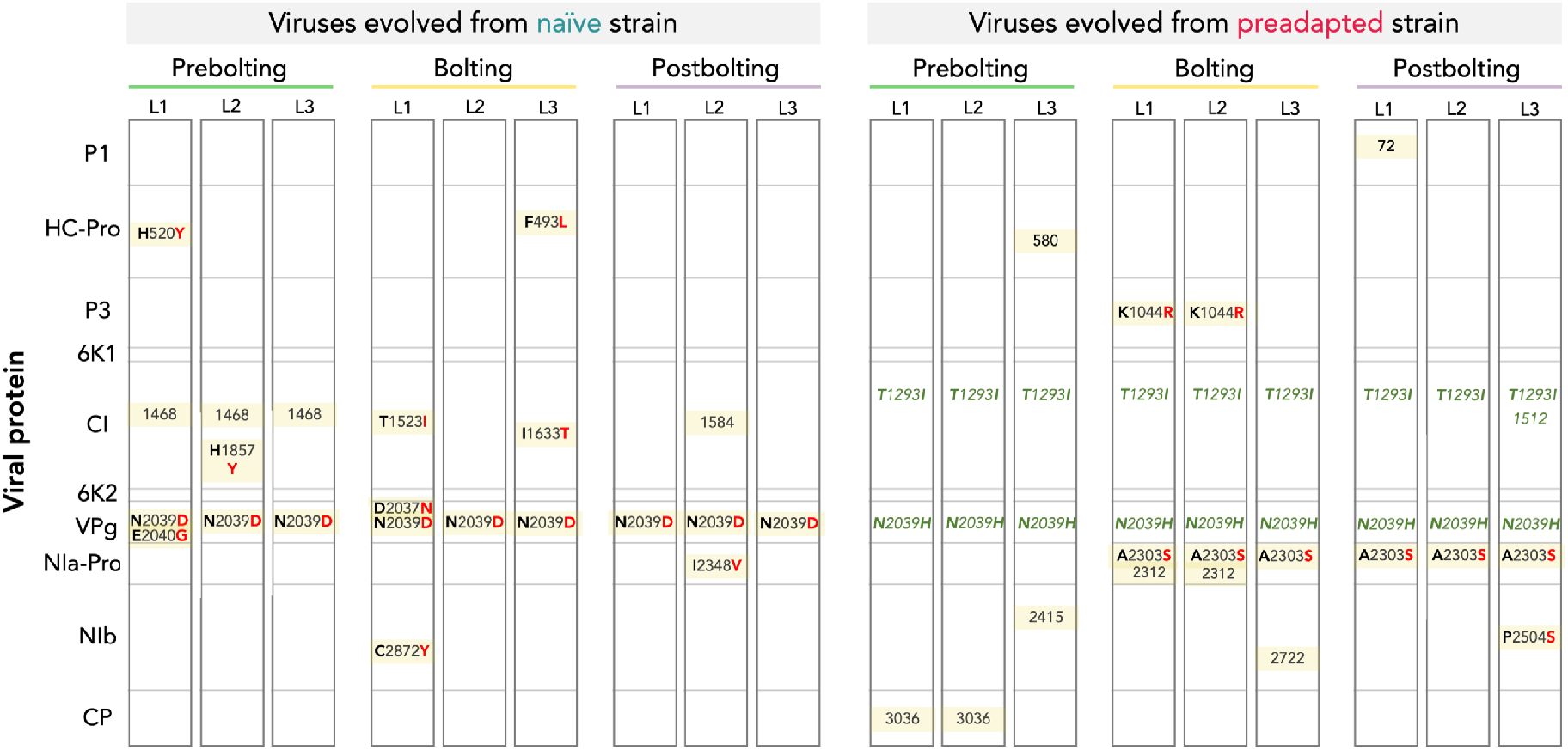
Mutations observed on every evolved lineage. The ancestral isolates, plant developmental stage and lineage are indicated above. Squares are proportional to the size of the cistron indicated in the first column. Synonymous mutations are indicated with the number of the nucleotide position in the polyprotein. Nonsynonymous mutations are indicated with the ancestral amino acid in black and the mutated one in red. Nonsynonymous mutations that were already present in the preadapted DV isolate are indicated in green.

For lineages evolved from the preadapted DV isolate, the mutation pattern was markedly different: no additional mutations were found in the *VPg* gene. For the three lineages evolved in prebolting plants, all the mutations were synonymous. By contrast, for lineages evolved in bolting and postbolting plants, all share nonsynonymous mutation NIa-Pro/A2030S (small non-polar by a polar change). In addition, two lineages evolved in bolting plants also shared mutation in P3/K1044R (conservative change of long positively charged side chains).

## Discussion

Host age constitutes an often-overlooked factor when studying plant-virus interactions and the outcome of an infection. In plants there are observations of the reassignment of resources between flowering and defense [27]. Classic life history evolution theories state that selection on any trait weakens as organisms age [50,51]. According to this assumption, fitness-related traits such as immune defence would be under weaker selection in older hosts; thus, a decline on host resistance to pathogens over time shall be expected. This assumption seems to hold for our pathosystem *A. thaliana* - TuMV.

Even though increasing resistance with age seems to be common in plants, the ARR defense mechanism that explains this phenomenon has mainly been described for plant-bacteria interactions [16-18]. Results with viruses are somehow contradictory in this regard. While there are examples of aged plants being more resistant to viral infections than juveniles, others show the opposite trend. García-Ruiz and Murphy [15] showed in the *Capsicum annuum* - CMV pathosystem that severity of symptoms and viral accumulation in non-inoculated leaves reached higher levels on juvenile than in mature plants. Levy and Lapidot [53] showed in the *Solanum lycopersicum* -tomato yellow leaf curl virus pathosystem that infected juvenile plants had significantly lower yield in comparation to mature ones. In sharp contrast yet in line with our observations, a recent study with the arabidopsis - tomato spotted wilt virus (TSWV) and arabidopsis - CMV pathosystems illustrates a developmentally regulated increase in susceptibility as plants become older under short day photoperiod [20]. Since arabidopsis is an annual plant that reproduces only once before dying, it makes sense that it adopts a strategy prioritizing reproduction over defenses.

The relationship between defense and host age may depend on the host and/or pathogen genetics and the interplays between both. Huang et al. [20] observed that developmentally regulated susceptibility does not occur in some host species such as tomato, pepper, or *N. benthamiana* (all *Solanaceae*) as it occurs for arabidopsis (a *Brassicaceae*). How specific or extended the age-dependent susceptibility is among other plant families calls for further studies. Huang et al. [20] results infecting arabidopsis with CMV and TSWV align well with our results infecting this plant with a set of viruses belonging to different genera: all of them infected better mature than juvenile plants. Whether this is a universal property of the way arabidopsis interacts with viral pathogens or just a spurious consequence of the chosen viruses and limited sample size (*n* = 11 viral species in total) needs to be further explored. Furthermore, to explore the effect of fitness differences among genetic variants of the same virus, we compared the results obtained for two isolates of TuMV with different past evolutionary histories. We observed that both isolates displayed the same host age dependence, infecting better mature hosts, although the room for phenotypic improvement for the poorly-adapted isolate was larger, as shown before for TuMV [47] and for bacteriophages used in phage therapy preadapted to *Pseudomonas aeruginosa* [54].

Our study also suggests, at least for TuMV, that prebolting plants are generally more susceptible to virus infection in short days than in long ones. This observation is in line with the results reported in [55], showing that reactive oxygen species (key elements for the successful activation of plant immune responses) on tobacco leaves kept in short days reached higher levels than when maintained under long days. More generally, it has also been described that photoperiod plays an important role in multiple phenotypes, including immunity [56]. However, the relationship between photoperiod and immunity may be pathosystem-dependent, as for example long day conditions increase susceptibility of sticklebacks to a parasitic flatworm [57]. Remarkably, we observed that the developmentally regulated susceptibility was independent of the host photoperiod.

Our results indicate that host-age can be an important factor on virus evolution. In our experimental pathosystem we have observed infection proceeds faster in mature host, indicating they are more susceptible. However, phenotypic rates of evolution are slower in the mature hosts than in the juvenile ones. This can be easily explained by differences in selective pressures: while mature hosts are permissive and represent a weak selective environment, juveniles are more restrictive to infection and represent a harsh selective environment. Thus, adaptive mutations in the latter might provide a disproportionally larger fitness benefit than in the former. From this, it is tempting to speculate whether certain developmental stages might select for viruses that specialize on infecting that particular stage or, by contrast, other more complex yet interesting possibilities exist: *e*.*g*., viruses selected in the more restrictive juveniles would behave as generalists infecting all age classes equally well whereas viruses selected in the more permissive mature plants would act as host age specialists. This test is now underway in the lab.

Depending on the developmental stage where plants were infected, the differential host-virus interactions caused viruses to face different evolutionary constraints that gave arise to different mutational spectra. Viral lineages evolved from the preadapted DV in bolting and postbolting hosts selected for the same mutation in the *NIa-Pro* cistron (A2303S). This suggests that this viral protein may have a key role during infection in these older stages but not in the juvenile one. Notably, DV-derived lineages evolved in prebolting juvenile plants show no fixed nonsynonymous mutations. This apparent genetic stability could be partially explained by the past evolutionary history of the DV isolate: since it was already preadapted for ten passages in prebolting conditions, we can confidently assume that specific adaptations to this developmental stage already took place. The presence of the same synonymous mutation in position 1468 of the *CI* cistron in all lineages evolved from the naïve AS virus in prebolting plants suggests that the selective pressure imposed by the host may be different depending on the host age. It is important to notice that certain selective pressures may still be independent of the host developmental stage, as it may be suggested by certain mutational events: (*i*) lineages evolved from the naïve virus in bolting and postbolting hosts have some different mutations in CI. (*ii*) Likewise, the ancestral DV isolate fixed mutations CI/T1293I during its original evolution in prebolting plants, suggesting that this protein may be targeted independently of the host age. This mutation has been retained in all DV-derived lineages irrespective of their host developmental stage. (*iii*) During its original evolution, DV also acquired mutation VPg/N2039H that has also been conserved in all its derived lineages. Interestingly, the lineages evolved from the naïve AS isolate had all fixed mutation VPg/N2039D in the same position. The selection on VPg mutants at position N2039 seems related with the function of this protein, as it is involved in virus movement, genome replication, and suppression of host antiviral RNA silencing [42]. Previous evolution experiments from our group had also described mutations in the same region of VPg on evolved viruses with increased virulence, independently of the arabidopsis genotype, its susceptibility to infection or the environmental conditions [58,59]. Similarly, mutations in this protein increased the fitness of TuMV in arabidopsis plants with knock-outs in the *eIF(iso)4E* and *eIF(iso)4G* genes [60] as well as the virulence of potato virus Y in resistant pepper plants [61].

This work was not intended to provide any insight into the molecular mechanisms behind the host-age dependent susceptibility to viral infection and its related consequences in virus evolution. Ongoing studies of the host transcriptomic and hormonal signaling responses to TuMV infection at each developmental stage will help us understand which host responses are responsible for the observed differences in susceptibility. A better understanding of the host response will also contribute to identify the selective pressures that viruses face during evolution in hosts of different ages.

## Conclusions

In this work, we aimed to study the impact of host age on plant’s susceptibility to viruses. We observed a positive correlation between host age and its susceptibility to multiple RNA viruses belonging to different genera. Focusing on a potyvirus species, TuMV, we have observed that this correlation occurs independently of the degree of adaptation of the viral isolate or the photoperiod the plant host is exposed to. We also studied the impact of host age on virus evolution. Firstly, we reported an increase on disease and symptomatology progression when comparing evolved viruses with their ancestral ones. This increase was more prominent the younger the host where the virus evolved was. Secondly, we show that host age conditions the spectra of mutations that appeared on the evolved viral lineages. Overall, this work confirms that plant aging may increases susceptibility to viruses and describes for the first time its impact on virus evolution. The results of this study will help to better understand the effect of host age on host-virus interactions and virus evolution.

## Acknowledgments

We thank Francisca de la Iglesia and Paula Agudo for excellent technical assistance. We also thank José A. Daròs, Pedro Gómez and Carmen Hernández for providing infected material for all viruses other than TuMV. This work was supported by grant PRE2020-094661 (Agencia Estatal de Investigación - FEDER) to I.M., BES-2016-077078 (Agencia Estatal de Investigación - FEDER) to R.G. and grants PID2019-103998GB-I00 (Agencia Estatal de Investigación - FEDER) and PROMETEU/2019/012 (Generalitat Valenciana) to S.F.E.

## Authors’ Contributions

I.M., data curation, formal analysis, investigation, visualization, writing-original draft. F.I., investigation, methodology, project administration. R.G., conceptualization, formal analysis, investigation, visualization, supervision, writing-original draft, writing-review and editing. S.F.E., conceptualization, formal analysis, funding acquisition, project administration, resources, supervision, writing-review and editing. All authors gave final approval for publication and agreed to be held accountable for the work performed therein.

